# CO_2_ emission at a hypersaline shallow lake at sediment-atmosphere interface. The significance of the organomineral upper crust as an active barrier

**DOI:** 10.64898/2026.01.27.701275

**Authors:** A. Butturini, O. Cabestrero, J. Ferriol, A. Blasco, M. Berlanga, P. Picart, Y. García de Fuentes, R. Gómez, J. Urmeneta, A. M. Romaní, Esther Sanz-Montero

## Abstract

Saline endorheic shallow playa-lakes are ecosystems susceptible to extreme geochemical changes because of severe hydro-climatic fluctuations. Under dry conditions, a rigid salt crust can separate the underlying sediments from the atmosphere. This interface is an organic-mineral assemblage of benthic biofilm encapsulated by evaporitic salts. It is well known that its structure, composition and consistency control water evaporation from underlying sediments, but its role in CO_2_ fluxes is unknown. We therefore measured the CO_2_ exchanges from sediments and the atmosphere in a hypersaline playa-lake characterized by a thin organic-mineral benthic crust upon drying. Results show that the largest CO_2_ release to the atmosphere occurs when ambient temperature and sediment humidity are high and low respectively. Nonetheless fluxes were lower than those reported for typical dry freshwater sediments and other hypersaline lagoons.

The dry crust contains sedimentary structures that likely reflect the gaseous pressure from the underlying sediments, and its removal provokes a significant increase in net CO_2_ fluxes.

Thus, this interface exerts physical control over both water evaporation and gases exchange such as CO_2_. Nevertheless, prolonged and severe droughts threaten the integrity of the crust: cracks together with bio-induced burrows and tunnels, likely create preferential pathways for CO_2_ leakage and enhance oxygen diffusion within the sediments and likely promote aerobic heterotrophic activity, explaining the CO_2_ leakage observed just below the organic-mineral crust.

## 1. Introduction

Hypersaline shallow lakes are endorheic inland ecosystems in arid, evaporite-rich regions that are sporadically flooded after severe rains. Upon flooding, a desiccation cycle begins; intense evaporation reduces water volume and salinity may rise by 35–45% (Butturini et al., 2022). Ultimately, the system dries, dissolved salts precipitate, and a rigid mineral crust can form at the surface, often persisting until the next major rainfall. Systems with this rigid interface are geomorphologically peculiar: the crust affects land erodibility, dust formation (Nield et al., 2015; Hamzehpou & Marcolli, 2024), soil albedo (Fujimaki et al., 2003), and water and heat fluxes by limiting evaporation of pore water in underlying sediments (Licsandru et al., 2022).

Recent studies on these systems focused on CO_2_ and CH_4_ fluxes across water–atmosphere or sediment–atmosphere interfaces to evaluate their contribution to the global carbon cycle (Cobo et al., 2024; Morant et al., 2024). These reports shows that hypersaline systems often (though not always) act as CO_2_ sinks when flooded and become net CO_2_ emitters when sediments are dry and exposed to the atmosphere (Appendix 1). Remarkably, little attention was given to sediment mineralogy, and most studies did not report whether a surface crust had formed. Little is known about links between CO_2_ fluxes and evaporitic mineral formation (Olson et al., 2023) or whether transformations of hydrated minerals affect gas exchange (Cabestrero et al., 2018). Consequently, it is logical to ask whether the upper crust influences CO_2_ exchange between the atmosphere and the sediments in hypersaline lakes.

According to these premises the present study had two main objectives:

1. To estimate the net CO_2_ exchange at the sediments-atmosphere interface in a hypersaline system under wet and dry conditions and thereby determine how water availability influences the magnitude and direction of CO_2_ fluxes.
2. To investigate the influence of the upper rigid salt crust on the modulation of CO_2_ fluxes.

This study focused on an endorheic hypersaline lake “La Muerte” (Monegros desert, Aragon, northeast Spain), whose bed is uniformly covered by a thin benthic biofilm. Upon drying, the biofilm transforms into a rigid organomineral layer (hereafter “OrgMinL”), where evaporitic minerals encapsulate the biofilm. Under desiccation, the OrgMinL hardens and cracks, forming polygonal patches with 30–50 cm segments.

To address the first objective, we measured in-situ CO_2_ fluxes at the sediment–atmosphere interface over one year to span a broad hydro-climatic range, though dry conditions predominated.

Further, a short controlled flooding experiment was performed in a small area under dry conditions to capture the impact of wet conditions. According to literature (Appendix 1) a net CO_2_ uptake is expected when sediments are water saturated whereas a release is expected when they are dry.

To meet the second objective, we combined in situ and laboratory measurements. In situ, CO_2_ fluxes were measured (a) before/after OrgMinL removal, (b) at multiple depths along a sediment profile, and (c) in zones with abundant OrgMinL fractures. In the lab, the focus was placed on assessing whether CO_2_ uptake by primary producers proliferating in the OrgMinL is significant. Concurrently, the petrography and sedimentary formations in the upper salt crust and underlying sediments were studied to identify structures potentially relevant for CO_2_ exchange.

If the OrgMinL limits evaporation from underlying sediments, it may also transiently retain CO_2_ released by the moist layer beneath; thus, CO_2_ escape should increase after OrgMinL removal. Under dry conditions, we further expect sedimentary structures in the upper salt crust to impede CO_2_ exchange, whereas fractures in the OrgMinL act as preferential degassing pathways. Consequently, under dryness, areas with dense salt-crust fracturing should emit more CO_2_ than intact surfaces. By contrast, in wet sediments, lower net CO_2_ release—or even net CO_2_ uptake—is expected owing to enhanced primary production in the upper layer.

## 2. Material & Methods

### 2.1 Study site

La Muerte lake (41°24’3.17”N; 0°15’39.16”O; Figure 1) is one of ∼100 shallow depressions in the Bujaraloz–Sástago endorheic complex, Southern Monegros Desert, Aragón (NE Spain), whose geomorphology and hydrology are detailed in Castañeda et al. (2013). The lake bed extends over 17.5 ha. Water is present 34% of the time, mainly between October and March, and is classified as “pristine” (Castañeda and Herrero, 2013).

**Figure 1.**
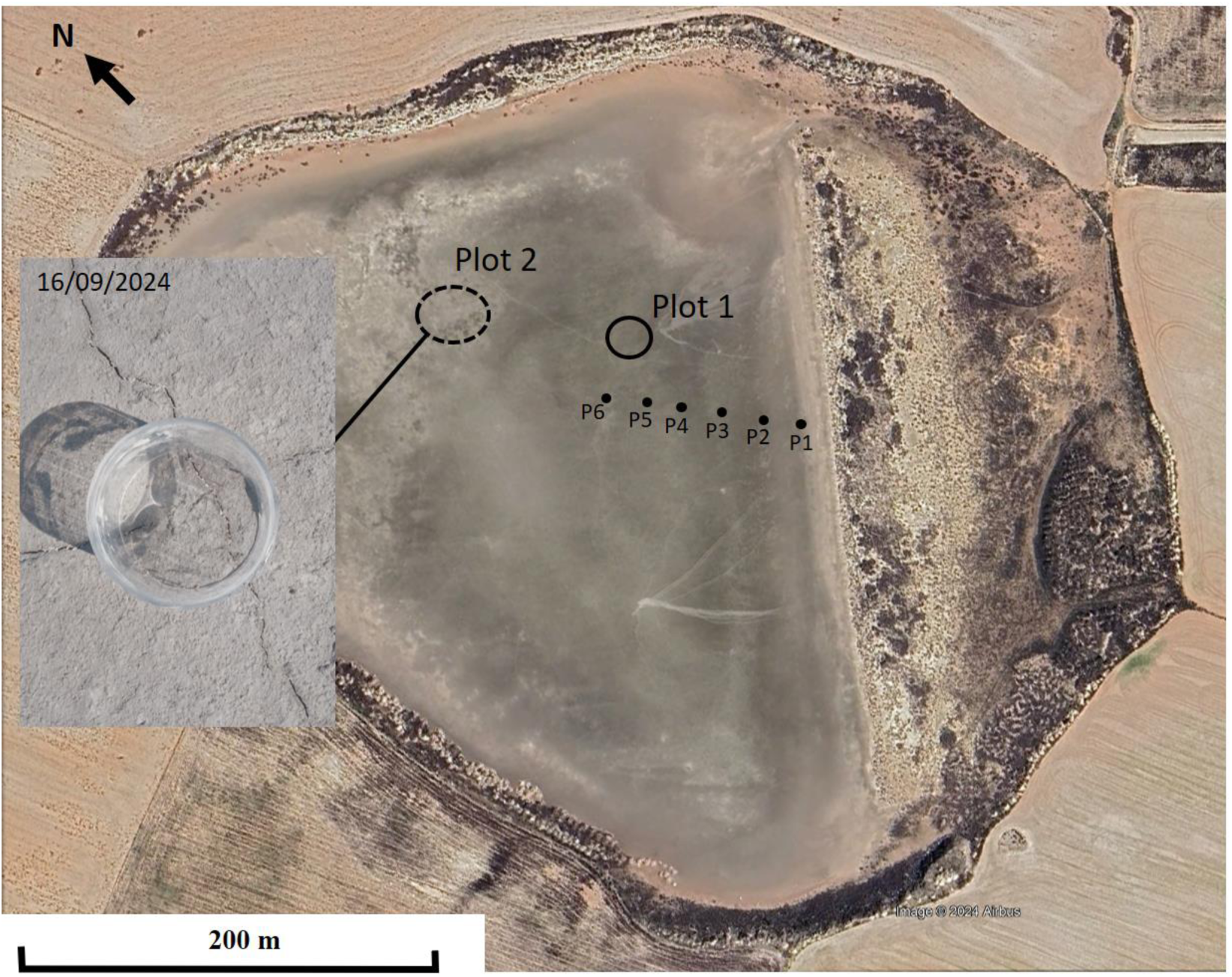
Aerial view of La Muerte lake. Satellite image acquired in March 2024 under dry conditions (Google Earth). The greyer area delimits the most humid portion of the lake bed. Plot 1: CO_2_ flux measured with/without the OrgMinL (June 2023–September 2024); sediments collected for mineralogical analysis. Plot 2: CO_2_ flux measured in intact and fractured OrgMinL under dry conditions (September 2024). Black dots indicate additional CO_2_ estimates along a longitudinal transect. Inset: detailed view of the dry lake surface in Plot 2. The cylinder has a basal area of 0.045 m^2^.

The bed is flat and uniformly covered by a thin monolayer (0.3–0.5 cm) of biofilm and minerals (Alcorlo et al., 1997). The upper 10 cm contain ∼15% organic matter. Detritic organic matter (mainly wind-transported woody fragments from surrounding shrublands) is ∼0.13±0.05 g m⁻².

Previous work revealed marked bio-structural and functional contrasts between the OrgMinL-bearing surface and underlying sediments (12–15 cm). Surface biomass (∼1 mgC gDW⁻¹) is nearly fivefold that of the subsurface, paralleling higher total protein, extracellular polymeric substances (EPS), prokaryote density, and chlorophyll; surface biomass is dominated by primary producers, whereas fungi are more prevalent below (Boadella et al., 2024).

The OrgMinL microbiome is dominated by Bacteria (82%), followed by Archaea (17%) and Eukarya (0.1%). Among bacteria, Pseudomonadota (31.85%), Bacteroidota (21.4%), and Cyanobacteria (18.4%) dominate; Euryarchaeota comprise 97% of Archaea. Twenty-six percent of the community is autotrophic and the remainder heterotrophic (Berlanga et al., 2024).

This study spanned June 2023–September 2024. During this period, partial flooding occurred only on January 29, 2024, when a water lamina <2 cm deep covered ∼20% of the surface, while the remaining sediments were water-saturated but unsubmerged. No surface water was detected on other dates except April 5, 2024, when a shallow pool (<1 m²) persisted within Plot 1. On June 29, 2023, and September 16, 2024, subsurface water-saturated sediments were observed in Plot 1. On all other sampling dates, the lake bed was dry, with no traces of water-saturated sediments.

### 2.2 Net CO_2_ fluxes at sediment–atmosphere interface

This study is based on in-situ and laboratory net CO_2_ flux measurements performed with intact sediment samples. The protocol adopted for measuring CO_2_ fluxes are described below.

### 2.2a) In-situ CO_2_ flux measurement

Measurements were performed:

1. At specific locations within two plots of La Muerte lake (“Plot 1” and “Plot 2”, Figure 1);
2. At six points aligned along a 100 m long transect (Figure 1) and;
3. At various sediment depths, from the surface to 20 cm depth at a specific location within Plot 1.

All in-situ CO_2_ flux measurements, except the depth profile (see below), were performed between 12:00 and 14:00 using a closed transparent plastic chamber (basal area: 0.015 m^2^ and height: 0.12 m) pushed 1 cm into the sediment.

### CO_2_ fluxes under dry and wet conditions

To address the first objective of this study, measurements were performed in Plot 1 on seven dates from June 2023 to September 2024. Moreover, on July 30^th^, 2024, under severe dry conditions and high ambient temperatures, a controlled in-situ flooding test was performed, during which a small area (0.2 m^2^) of dry bed sediments was flooded for 2 hours. Net CO_2_ fluxes were measured before and after flooding to quantify the impact of an abrupt rewetting of sediments on these fluxes.

To explore the relevance of spatial heterogeneity on net CO_2_ fluxes, six additional measurements were performed on four dates from July 2023 to March 2024 along a 100 m long transect extending from the outer limit of the lake and ending near Plot 1 (sites P1 to P6 in Figure 1).

### Effect of the OrgMinL on CO_2_ fluxes

To address the second objective of this study, we compared the net CO_2_ flux estimations obtained at a location within Plot 1 with preserved sediments to those obtained immediately after the removal of the upper OrgMinL. These comparisons were performed seven times on four dates from June 2023 to July 2024. The OrgMinL was removed manually with a spatula.

To analyze the effect of the presence of fractures in the OrgMinL on net CO_2_ fluxes, we selected Plot 2, because it was drier and more fractured than Plot 1. Twelve CO_2_ flux measurements were performed in this plot under dry conditions on September 16^th^, 2024. Six measurements were made at locations with fractures, while the remaining six were made at adjacent locations without any visible fractures. At the selected locations, the “Fracture density” (F_d_) was defined as the ratio between the area occupied by fractures to the total area of the crust being incubated in the chamber.

To further investigate the relevance of the OrgMinL and underlying sediments on CO_2_ fluxes, a specific sediment depth profile was performed in Plot 1 under dry conditions, on July 30^th^, 2024. Net CO_2_ flux measurements were performed at seven sediment depths: 0 (i.e. intact OrgMinL), 0.3 cm (without the OrgMinL) 1, 3.5, 8, 14 and 20 cm. Measurements were made by pushing a small closed transparent plastic cylinder (basal area: 0.0015 m^2^ and height: 0.04 m) into the sediment. To perform these measurements, the sediment was removed gradually by hand with a spatula to the target depth. Sediment samples from each depth were collected and stored at 4 °C in order to estimate the water activity (a_w_), water gravimetric content (W_c_), AFDW (ash free dry weight, a proxy of organic matter content), prokaryote density (ProkD), and β-glucosidase extracellular enzyme activity (B-Glu) in the laboratory (see details below).

### 2.2b) Ex-situ CO_2_ flux measurements. Light-dark incubations

Focusing on the potential effect of photosynthetic primary production in the OrgMinL (Boadella et al., 2024; Berlanga et al., 2024) on the net CO_2_ fluxes, the following three ex-situ experiments (labelled Ex-Situ Exp#1, 2 and 3 respectively) were performed in the laboratory.

Ex-Situ Exp#1 evaluated the impact of primary producers on CO_2_ fluxes in samples containing both the OrgMinL, where photosynthetic organisms develop, and the underlying 4–7 cm of sediments. This test most closely mimicked field conditions. In contrast, in Ex-Situ Exp#2 and Exp#3, measurements were performed with samples composed exclusively the OrgMinL previously detached from the sediments. These two experiments specifically aim to quantify primary production in the OrgMinL by minimizing interference from CO_2_ released from the underlying sediments.

These three experiments are described in detail below.

In September 2024, under dry conditions and in water-unsaturated sediments, four undisturbed sediment “bricks” (20×14×7 cm), from Plot 1, were collected and transported to the laboratory. The thickness of the samples ranged between 4 and 7 cm. Three samples were preserved in transparent sealed plastic chambers to maintain moisture levels, whereas the fourth was stored in an identical container but left open to dry further.

Ex-Situ Exp#1): Within a week after sampling, each undisturbed sample was incubated at ambient temperature under natural light and subsequently under dark conditions. Both incubations were performed in triplicate for 15 minutes each and were conducted in the same chambers used at the time of sampling. Measurements started after a 30-minute light/dark adaptation period and were performed from 13:00 to 15:00, with light radiation oscillating between 35 and 50 Klux.

Ex-Situ Exp#2): Next, three subsamples (of known area, 7–18 cm²) of the OrgMinL were carefully detached from each of the three moist samples used in the previous experiment. These nine OrgMinL subsamples were placed in small transparent sealed plastic cylinders (radius: 0.0015 m²; height: 0.04 m) and incubated for 15 minutes, during which they were exposed alternately to natural light and darkness, under the same conditions described in Ex-Situ Exp#1.

Ex-Situ Exp#3): Once Ex-Situ Exp#2 was complete, four of the nine OrgMinL subsamples were submerged in aerated water from La Muerte (previously filtered through 0.22 μm nylon filters) for 1 week at 25 °C under natural light-dark conditions. A preliminary test indicated that a week in water and natural light-night conditions is sufficient to boost the growth of cyanobacterial filaments on the surface of the OrgMinL (Appendix 2). After rewetting, the samples were incubated for 15 minutes under natural light, followed by incubation in darkness, under the same conditions described for Ex-Situ Exp#1 and #2, after a 30-minute adaptation period.

In all experiments, light intensity was measured using a HI97500 Luxmeter (Hanna).

### 2.2c CO_2_ incubation protocol and flux calculations for in-situ and ex-situ measurements

In the in-situ measurements, a transparent closed chamber was plugged ∼1 cm into the sediments. The chamber was equipped with an NDIR EZO-CO2 sensor (Atlas Scientific Inc.), a LuminOx optical oxygen sensor (SST Sensing Ltd) that integrates temperature and atmospheric pressure sensors, and a DHT11 humidity+temperature sensor (DFRobot). All sensors were connected to an Arduino DUE that communicated with a portable PC using CoolTerm. Before each measurement, the EZO-CO2 sensor was calibrated in the lab with a calibrated CO2 gas analyzer IRGA EGM-4, together with a two-point calibration using pure N2 (0 ppm CO2) and a 3970 ppm CO_2_ standard. Each incubation lasted ∼15 min; data were stored every 2 s. CO_2_ readings are in μmol/mol. To estimate the net CO2 flux (FCO2) we first estimated the volume V(T,P) (L) of one mole of gas according to ideal gas law:

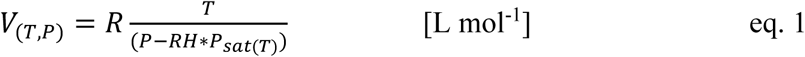

where R is the ideal gas constant 0.082 atm L mol^-1^ K^-1^, T is the air temperature during the incubation (K), P is the air pressure (atm), P_sat(T)_ is the saturation water vapour pressure at temperature T estimated using the Antoine equation, and RH is the relative humidity.

Then, each CO_2_ sensor reading “a” was converted into CO_2_ mass per unit of area of the chamber (A, m^2^) according to the following formula:

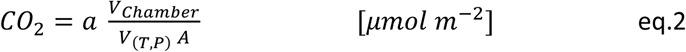

Where V_chamber_ is the volume of the head space. Finally, CO_2_ flux during each incubation was estimated assuming a linear change of CO_2_ mass over time:

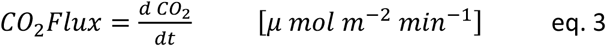

For Ex-Situ eperiments all chambers were equipped with a NDIR EZO-CO_2_ and a DHT11 humidity+temperature sensor (DFRobot), both connected to an Arduino DUE. Data were recorded every 2 sec. Net CO_2_ fluxes were calculated as described above.

### 2.3 Sediment relative humidity, water activity and organic matter content

Sediment relative soil mositure (RH) in Plots 1 and 2 was recorded on each sampling date. RH was measured using a HP23-AW-A (Rotronic Instrument Corp.) equipped with a HC2 humidity and temperature probe. The sensor was plugged into the sediment to a depth of 3.5 cm and humidity was recorded after the sensor had been measuring for 3 hours.

Water activity (a_w_), water content (Wc, %), and organic matter were measured in the in-situ sediment profile (Plot 1). Water activity was measured with an HP23-AW-A sensor (Rotronic Instrument Corp.), with 40-min readings to ensure stability; aw was also measured in sediment and OrgMinL samples used for ex-situ experiments. Sediment Wc was calculated as wet minus dry mass; samples were dried at 60 °C for 72 h before weighing. For organic matter, the same dried samples were combusted for 4.5 h at 450 °C (MF12-124, Hobersal, Barcelona, Spain) and re-weighed. Results are expressed as grams of ash-free dry weight (AFDW) per gram of dry weight (DW).

### 2.4 Prokaryote density and β-glucosidase enzyme activity

Sediment subsamples (1 cm³) from the in-situ profile at Plot 1 were collected to quantify prokaryote density and β-glucosidase (B-Glu) extracellular activity. Subsamples for prokaryote density (ProkD) were stored at −20 °C and analyzed one week later by flow cytometry (FACSCalibur, Becton Dickinson) following Boadella et al. (2024). β-glucosidase enzyme activity (b-Glu) is a proxy of the potential of heterotrophs to degrade labile organic matter sources into simple polysaccharides (i.e. decomposition of cellobiose or small oligomers to obtain glucose). β-Glu was estimated in fresh samples (1 cm^3^) according to the protocol detailed in Boadella et al. (2024).

### 2.5 Chlorophyll concentration

Chlorophyll was measured in the OrgMinL on January 29, 2024 (partially flooded) and July 30, 2024 (dry): 13 samples per date (10–20 cm²), plus all ex-situ samples. Pigments were extracted in 90% acetone for 24 h in darkness, sonicated 10 min, and filtered through GF/F filters. Spectra (300–1000 nm; 1-cm path) were recorded (Shimadzu UV-1700). Chlorophyll-a was calculated using Ritchie’s spectrophotometric models (2006), assuming cyanobacterial dominance (Berlanga et al., 2024). The 430/665-nm absorbance ratio (Margalef index) was used as a proxy of ecological status of chlorophyll: high values indicate a high relative proportion of carotenoids versus chlorophyll which indicates stressed photoautotrophs.

### 2.6 Petrography

Samples comprising the OrgMinL and the upper 1 cm of underlying sediment were collected at Plot 1 in bags (1–2 kg) and plastic containers (200 g). Thin sections were prepared with ethanol (96%) to prevent mineral dissolution and were subjected to freeze-drying to maintain textural features. OrgMinL was consolidated by resin impregnation. Optical analyses of bulk samples and thin sections were conducted using a Leica MZ16 lens and an Olympus BX51 optical microscope. Porosity was assessed at different scales by mapping photographs of resin-impregnated plugs and thin sections (Marín et al., 2021).

### 2.5 Data analysis

Linear regressions assessed how net CO_2_ fluxes in Plot 1 varied with air temperature, sediment humidity, and their joint effects; the optimal model was the one with the lowest Bayesian Information Criterion (BIC; Schwarz, 1978). Significance was set at p < 0.05 throughout. Paired t-tests compared net fluxes at the same locations in Plot 1 (n = 7) with and without the OrgMinL. A second paired t-test, in Plot 2, contrasted measurements from sites (n = 6) with versus without OrgMinL fractures.

A multiple linear model (MLM) examined the relationship between net CO_2_ fluxes and the six predictors measured along the sediment profile (T, a_w_, W_c_, AFDW, b-Glu and ProkD) at discrete horizons down to 20 cm. Because sealed chambers integrate CO_2_ production across the entire profile, not just surface sediments, each flux estimate at a given depth was related to a depth-weighted average of predictor values calculated across the full sediment column. Therefore, each measurement of CO_2_ at a given depth is associated with the weighted average of the six predictors measured at the different depths in the sediment column.

To identify the optimal MLM model we adopted the following strategy:

First, we defined *L_n_*, the list, of the six potential descriptors:

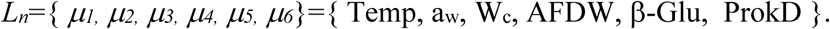

Then, all i distinct proper subsets of *L_n_* were extracted:

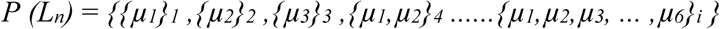

where *i*=2^n^-1. In this case n=6, there were a total of 63 subsets of *Ln*. However, subsets containing more than five variables were excluded due to the limited sample size of seven observations, which limits the maximum number of predictors in an MLM to five. Therefore, the real number of subsets was *i*=62.

Subsequently, an MLM was run for each *i* subset. Among all possible *i* models, the optimal one is that with the lowest BIC value (Schwarz, 1978).

## 4) Results

### 4a) Sediment humidity, water activity and organic matter content

Under dry conditions, the moisture content of the upper 3 cm ranged from 78.5% to 90.1%, with the lowest values on December 1^st^, 2023 and June 29^th^, 2023, and the highest on January 29, 2024. Figure 2 shows vertical profiles of water percentage (panel a), water activity a_w_ (panel b), and AFDW (panel c) in the upper 20 cm of lake-bed sediments under dry conditions (July 30^th^, 2024 sample). Both water percentage and a_w_ increased with depth, with the sharpest shifts at the interface between the OrgMinL and subjacent sediments. Notably, the OrgMinL combined relatively high water content with the lowest a_w_. Concurrently, the OrgMinL had the lowest AFDW, whereas the sediments immediately beneath it exhibited the highest organic-matter accumulation.

**Figure 2.**
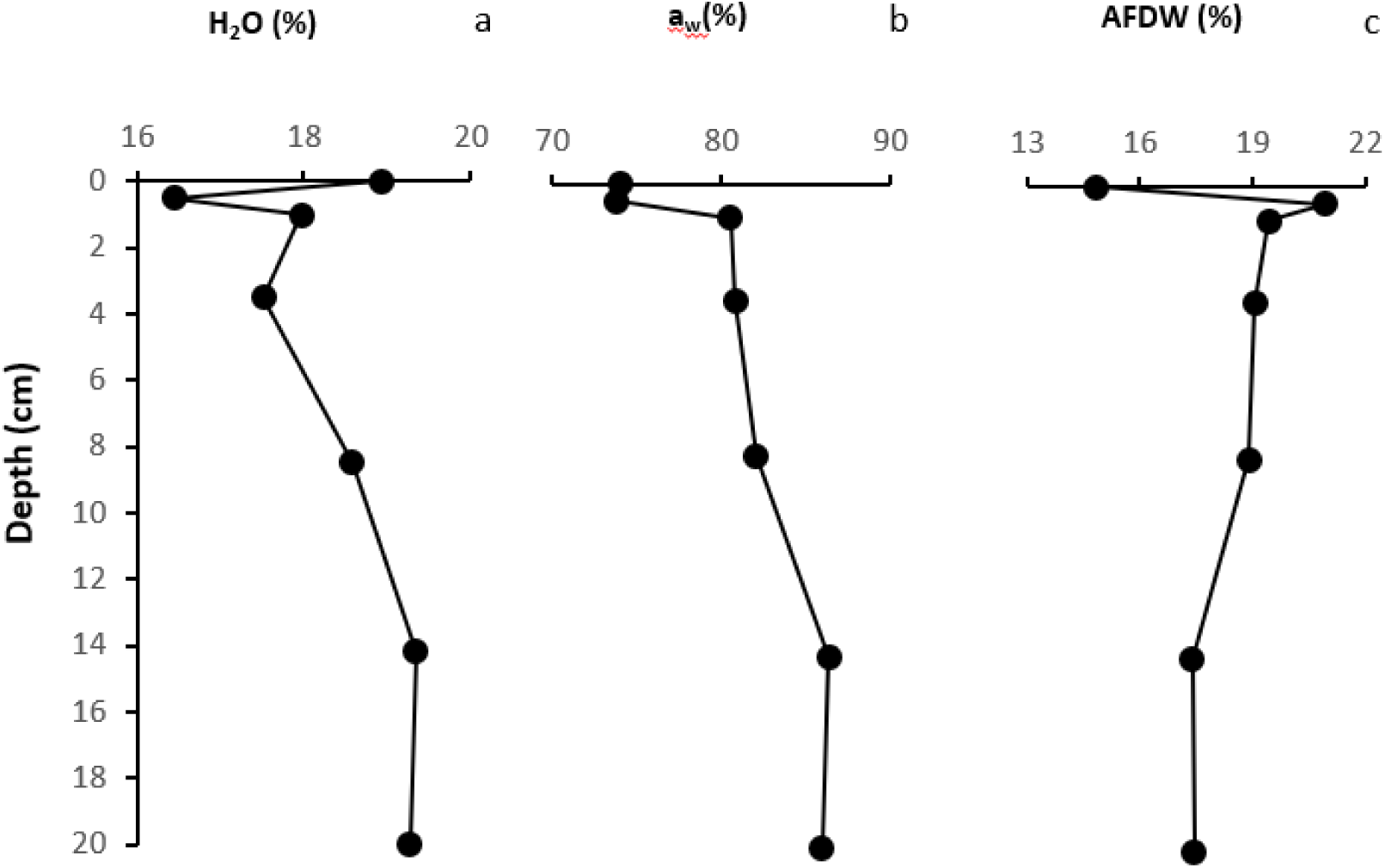
Vertical profiles of water content (panel a), water activity (b) and organic matter content (c) recorded in a 20-cm deep sediment profile. Measurements were performed under dry conditions in July 2024.

### 4b) Structure and composition of the sediments

The upper layer, roughly 0.5 cm thick, contained sedimentary structures that resulted from the initial growth of microbial mats under wet conditions, and subsequently, following their degradation, hardening after drying and mineral precipitation. Features include small domes, blisters (to 3–4 cm long, 1 cm high), and folds (to 10 cm long), locally detached from the substrate to create voids between the OrgMinL and sediments (Figure 3D and I), which became evident upon collapse (Figure 3F). An irregular polygonal crack network with upturned margins overprints these structures (Figure 3G, Appendix 3), fragmenting the organomineral crust (Figure 3B, D, E). The OrgMinL and underlying sediments were further disrupted by branched burrows up to 1 cm long and by tunnels, commonly unroofed (Figure 3A, E), and by piles of peloids from arthropod excavation (Figure 3E). Additional indicators of mat destruction include ripple marks, deer footprints impressed into the sediment (Figure 3H), and elongate drag marks left by small rocks. The OrgMinL overlied 1–2 cm thick layers deposited in successive episodes of sedimentation (Figure 3A). The underlaying sediments comprised intrasediment-grown lenticular to prismatic gypsum crystals (0.1–1 mm in size; Figure 3B) and clastic silt-sized quartz and feldspars sediments, whereas the OrgMinL consisted of carbonaceous microbial laminae and mineral precipitates. Locally, the organic laminae were folded and firmly attached to the bed, evidencing their elasticity and cohesiveness (Figure 3B, D), but more commonly the layer was degraded, with clots and irregular pores (Figure 3C). The clots consisted of mixtures of precipitated minerals and decayed cyanobacterial filaments with their EPS; cubic halite and acicular or tabular sulfates—mostly bloedite—were the main phases precipitated around the degraded remains (Figure 3C). Calcite and dolomite also precipitated in pores around the degraded filaments (Figure 3C). Pores in the OrgMinL, generated by organic decay and bioturbation (burrows) tended to be vertically and horizontally elongated (Figure 3A–C; Appendix 3). Burrows showed elongated and rounded sections that sometimes contained remains of the burrowers (Appendix 3) and halite crystals (Figure 3I–J), though most were empty or host-sediment filled. Total porosity from thin-section microphotographs ranges from 9.32% in the most compact crustal sediment to 33.71% in the degraded mat, with pore sizes of 10–400 μm in both cases. Porosity in the impregnated plug is 11.49%, with pore sizes from 50 μm to 1.5 cm (Appendix 4).

**Figure 3.**
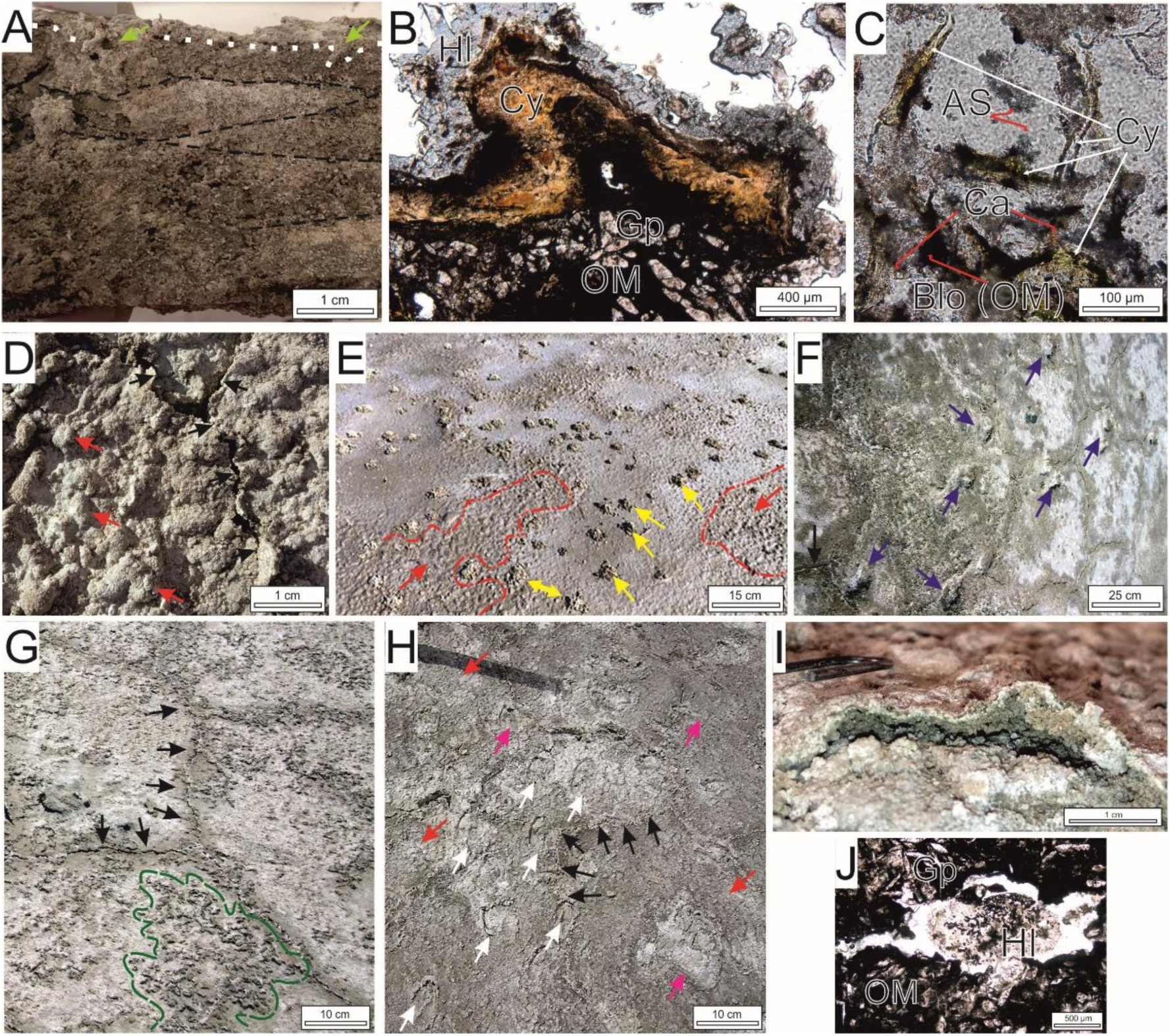
Structures in the OrgMinL coating the surface of the dry lake. A) Sediment profile showing a salt-encrusted microbial mat (dotted white line) with filled burrows (olive arrows), overlying strata of previous episodes (dashed black lines). B) Folded microbial carbonaceous laminae (Cy) coating intrasediment-grown gypsum (Gp); surface encrusted with halite (Hl). C) Porous layer with degraded cyanobacterial filaments (Cy) and EPS (OM), encrusted by sulfate acicules (AS), bloedite (Blo), and carbonate microcrystals (Ca). D) Elevated blisters (red arrows) amid cracked, partially detached OrgMinL (black arrows). E) Blistered morphologies (red arrows; red dashed line) and piles of peloids from beetle/wasp excavation (yellow arrow). F) Depressions from cracking and collapse of gas domes (dark blue arrows); black arrow marks mineral precipitates in a desiccation crack. G) Irregular polygonal cracks (black arrows) and a network of branched burrows beneath a thin microbial mat (green dashed area). H) Deer tracks (white arrows) on a cracked bed (black arrows) disrupting blistered (red arrows) and pitted features (pink arrows). I) Cavity within a blister. J) Elongated pores with halite crystals (Hl) cementing the host sediment dominated by gypsum (Gp) and decomposing organic matter (OM). Note that B, C, I, J are polarized-light microphotographs.

### 4d) CO_2_ fluxes at salt crust–atmosphere interface

In Plot 1, the CO_2_ net fluxes ranged between –29.5 and 32 μmol/m^2^/min. Conversely, maximum uptakes coincided with the coldest, most humid period, under water-saturated sediments (Figure 4), when chlorophyll-a rose to 13±3.5 μg cm^-2^ and the Margalef index declined to 2.57±0.16.

**Figure 4.**
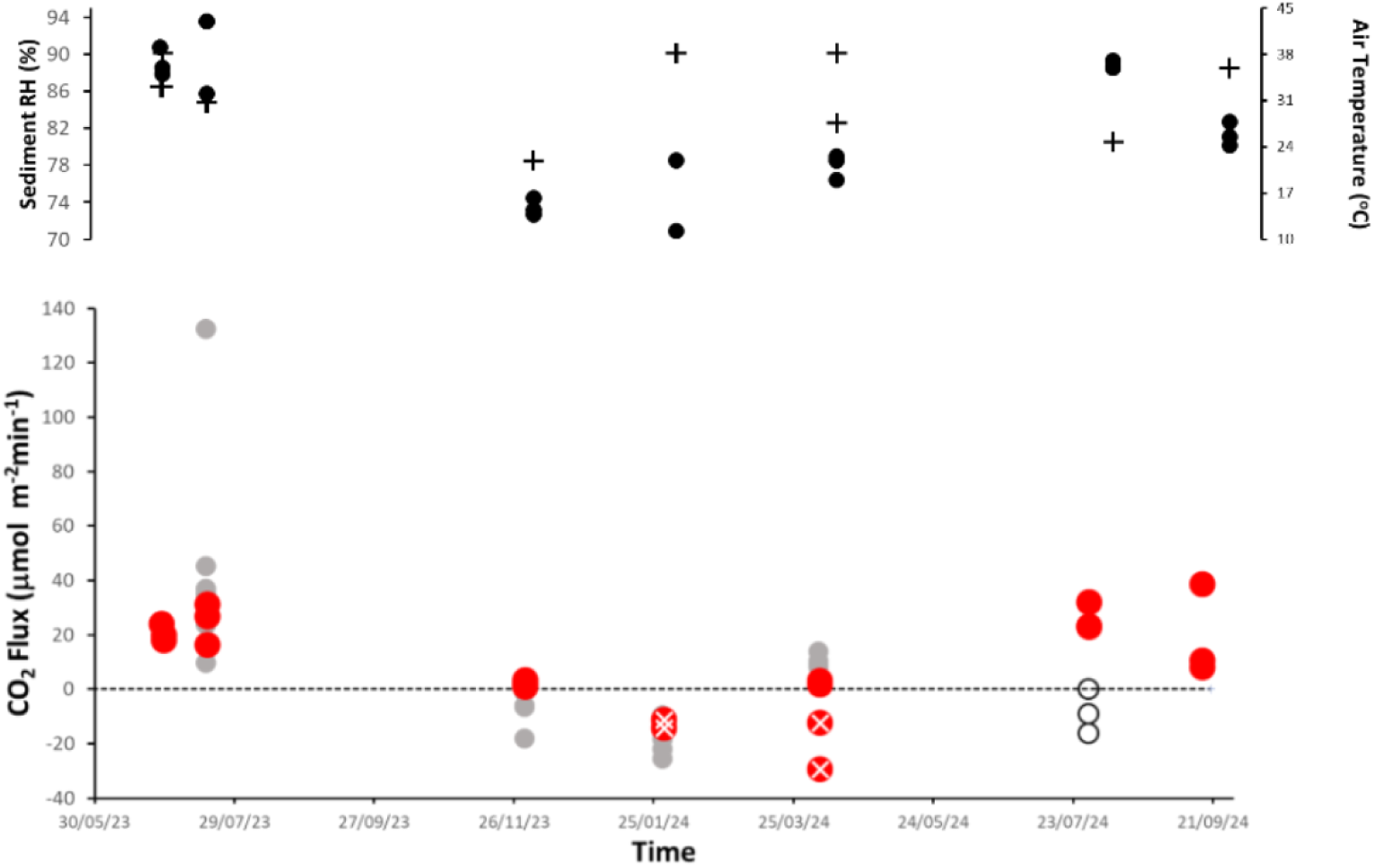
Temporal dynamics of net CO2 fluxes at the salt-crust–atmosphere interface in Plot 1 during the study period (red dots). Water-saturated sediments are marked by a white cross. Grey dots denote CO2 fluxes along the longitudinal transect. White circles indicate CO2 fluxes in Plot 1 after a 2-h controlled inundation. Positive and negative values denote net emission and consumption, respectively. Black dots show air temperature; black crosses, sediment relative humidity (RH%). On two dates, two RH% values are shown because fluxes were measured at two locations with different RH%.

In contrast, the highest net CO_2_ uptakes occurred during the coldest and most humid period when sediments were water-saturated (Figure 4) and chlorophyll-a increased up to 13±3.5 μg cm^-2^ and the Margalef index decreased to 2.57±0.16. Accordingly, CO_2_ net fluxes were significantly and positively related to temperature (r^2^=0.55, p<0.05, BIC=171; Figure 5). Net CO_2_ fluxes were not significantly related to sediment moisture (r^2^=0.07, p>0.05). However, combining temperature and sediment mositure generated a linear model (r^2^=0.647, p<0.05, BIC=169.5) that was a little more robust than the model that included temperature alone.

**Figure 5.**
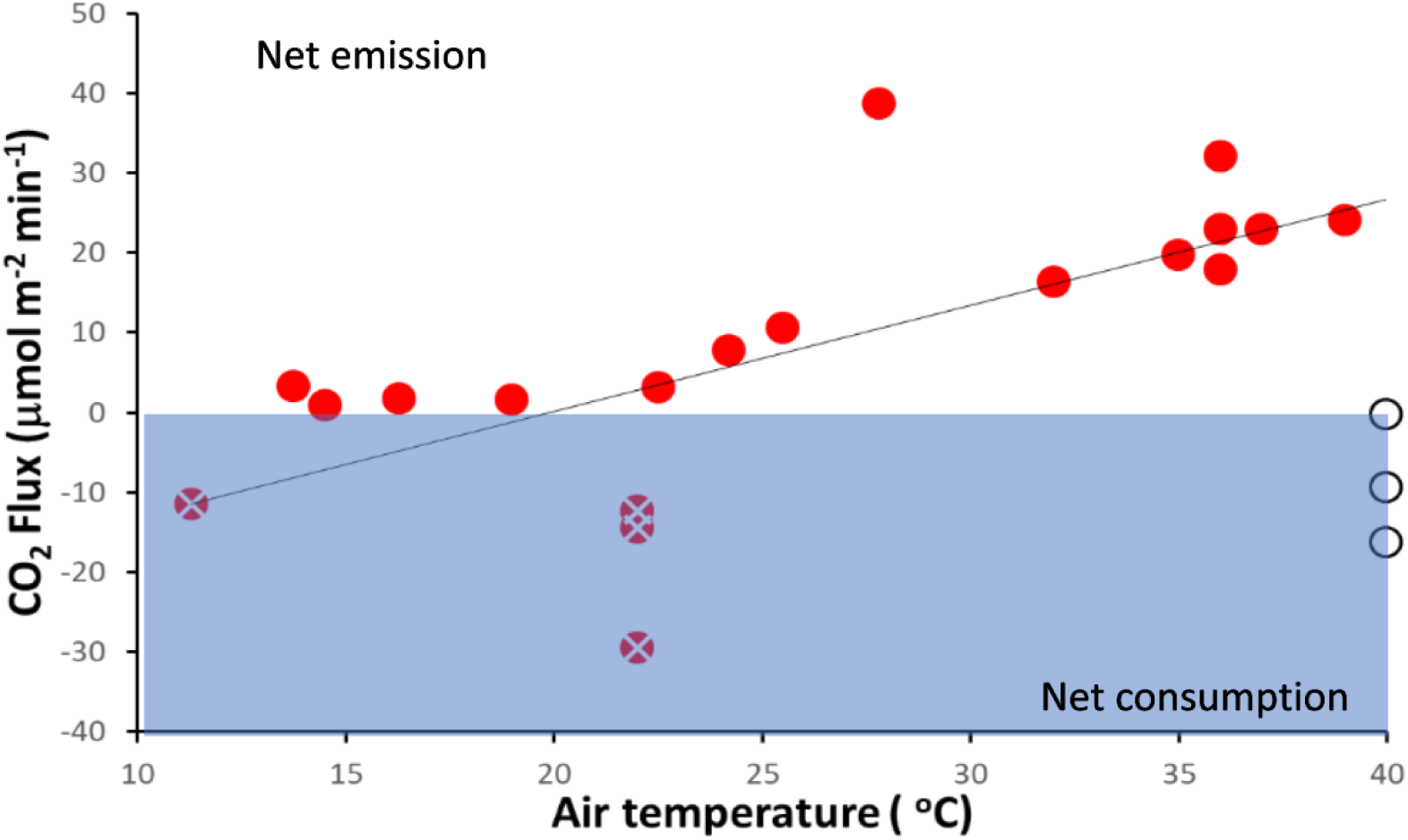
Relationship between CO_2_ fluxes measured at the salt-crust–atmosphere interface in Plot 1 and air temperature. Symbols are the same as in Figure 4. Solid line shows the regression model (r^2^=0.55, p<0.05). White circles (controlled inundation test) were not included in the regression model. Blue shading indicates negative net CO_2_ fluxes (i.e. net CO_2_ fixation).

The effect of water availability on net CO_2_ flux magnitude was further shown by a controlled in-situ flooding on July 30th, 2024, when air temperatures reached up to 40 °C. After 2 h of flooding, net CO_2_ emission reversed to a weak net CO_2_ uptake (Figure 5).

Net CO_2_ flux estimates along the longitudinal transect matched those in Plot 1 and mirrored the same temporal pattern (Figure 4). No spatial patterns in CO_2_ fluxes were detected, except on the July 2023 date, when the lowest and highest CO_2_ emission fluxes occurred at the most peripheral and innermost points, respectively.

### 4e) CO_2_ fluxes after removal of the OrgMinL

In all seven replicates, removal of the salt crust led to an increase in CO_2_ emission compared to the fluxes observed just prior to removal in the intact sediment (t_paired_ =2.55, d.f.,=5, p<0.05). The increase ranged from 26% to 140% (Figure 6).

**Figure 6.**
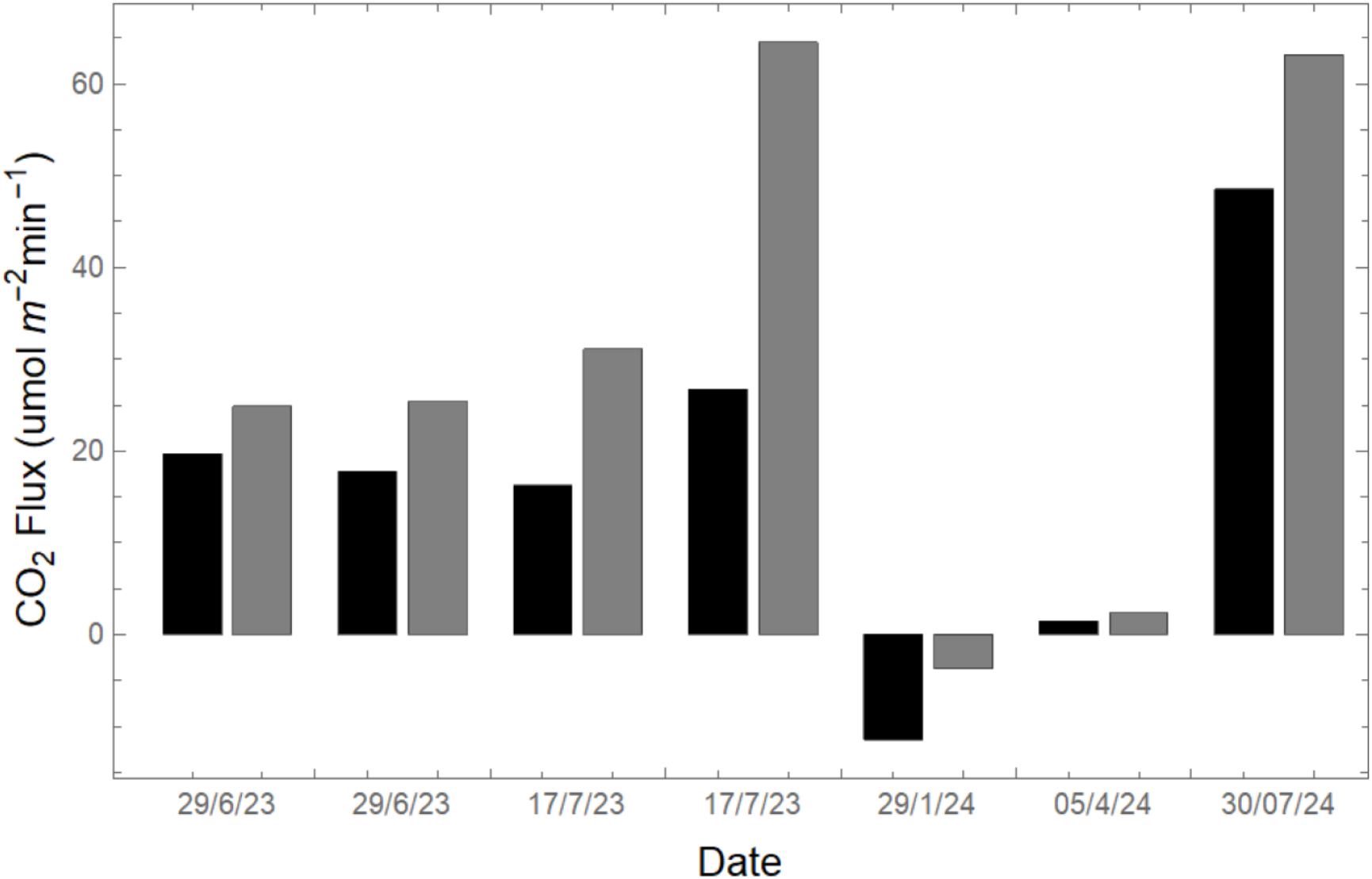
In-situ CO_2_ fluxes with (black bars) and without the upper OrgMinL (grey bars).

### 4f) CO_2_ fluxes at different sediment depths

On July 30^th^, 2024, under dry conditions, in-situ CO_2_ fluxes were estimated at different depths in sediment up to 20 cm below the OrgMinL. In this test, removal of the upper OrgMinL induced an increase in net CO_2_ flux as similarly reported in the previous section, the largest net CO_2_ flux being observed at z=3.5 cm. Below this depth, the CO_2_ flux decreased progressively with depth, revealing that most CO_2_ flux originates within the upper 5–8 cm of sediments (Figure 7). Accordingly, ProkD and β-glucosidase activity also decreased with depth. After evaluating all possible MLM models, the optimal model (with the lowest BIC value) revealed that CO_2_ flux along the sediment profile was positively related to AFDW and a_w_ (r^2^=0.8, p<0.05, BIC=1.8), with the latter variable being the most significant (**Appendix 5**).

**Figure 7.**
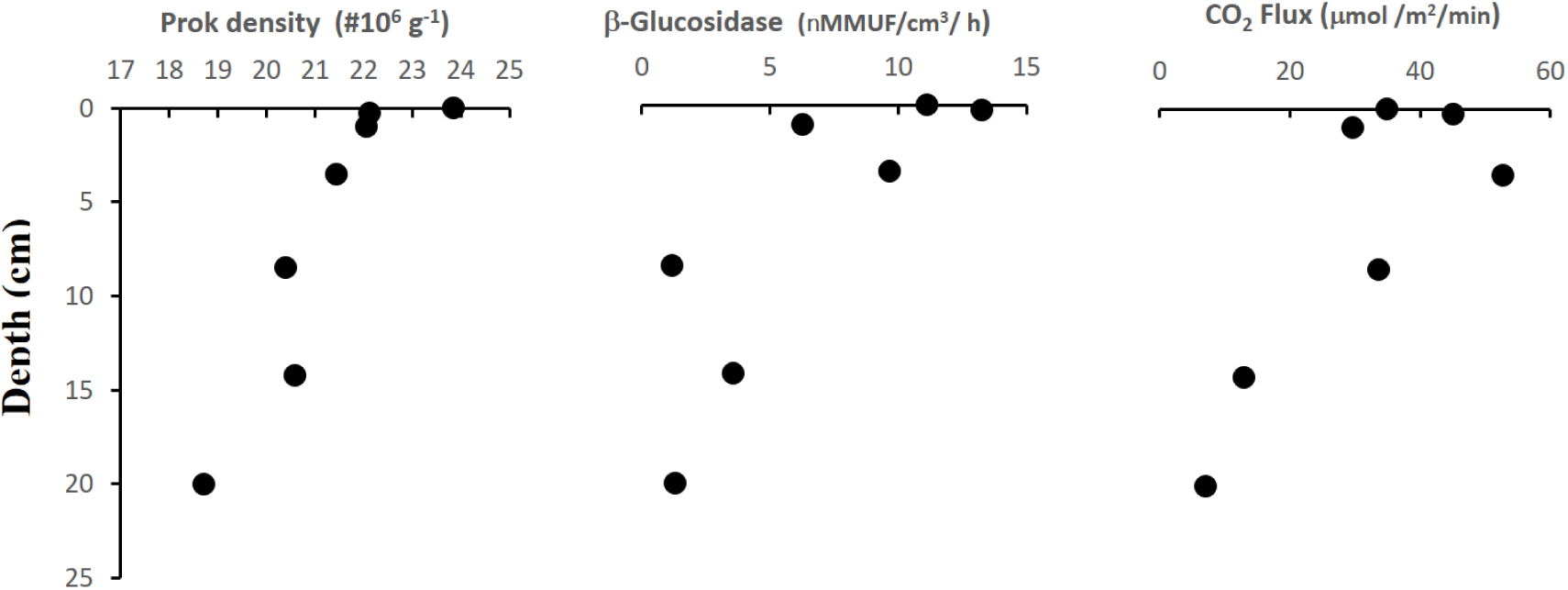
Vertical profiles of prokaryote density, β-glucosidase extracellular enzyme activity and net CO_2_ flux in the upper 20 cm of sediments. These measurements were performed in a sediment profile collected under dry conditions in July 2024. Figure 7. Relationship between CO_2_ fluxes and temperature reported in different hypersaline systems (dots) and dry freshwater systems (squares). CO_2_ fluxes reported in this plot were obtained using the closed chamber method. Black dots: this study; red dots: a hypersaline lake (Bourhane et al., 2023); orange dots: hypersaline tidal flats (Brown et al., 2021); grey dots: exposed Great Salt Lake bed sediments (Cobo et al., 2024); blue dots: hypersaline lake (Thomas et al., 2022); grey squares: exposed dry sediments from freshwater ecosystems (Keller et al., 2020). The range of “y” axes is constrained in the –50 to 200 μmol CO^2^ m^-2^ min^-1^ interval. Appendix 9 shows the same plot but with the full “y” range.

**Figure 7.**
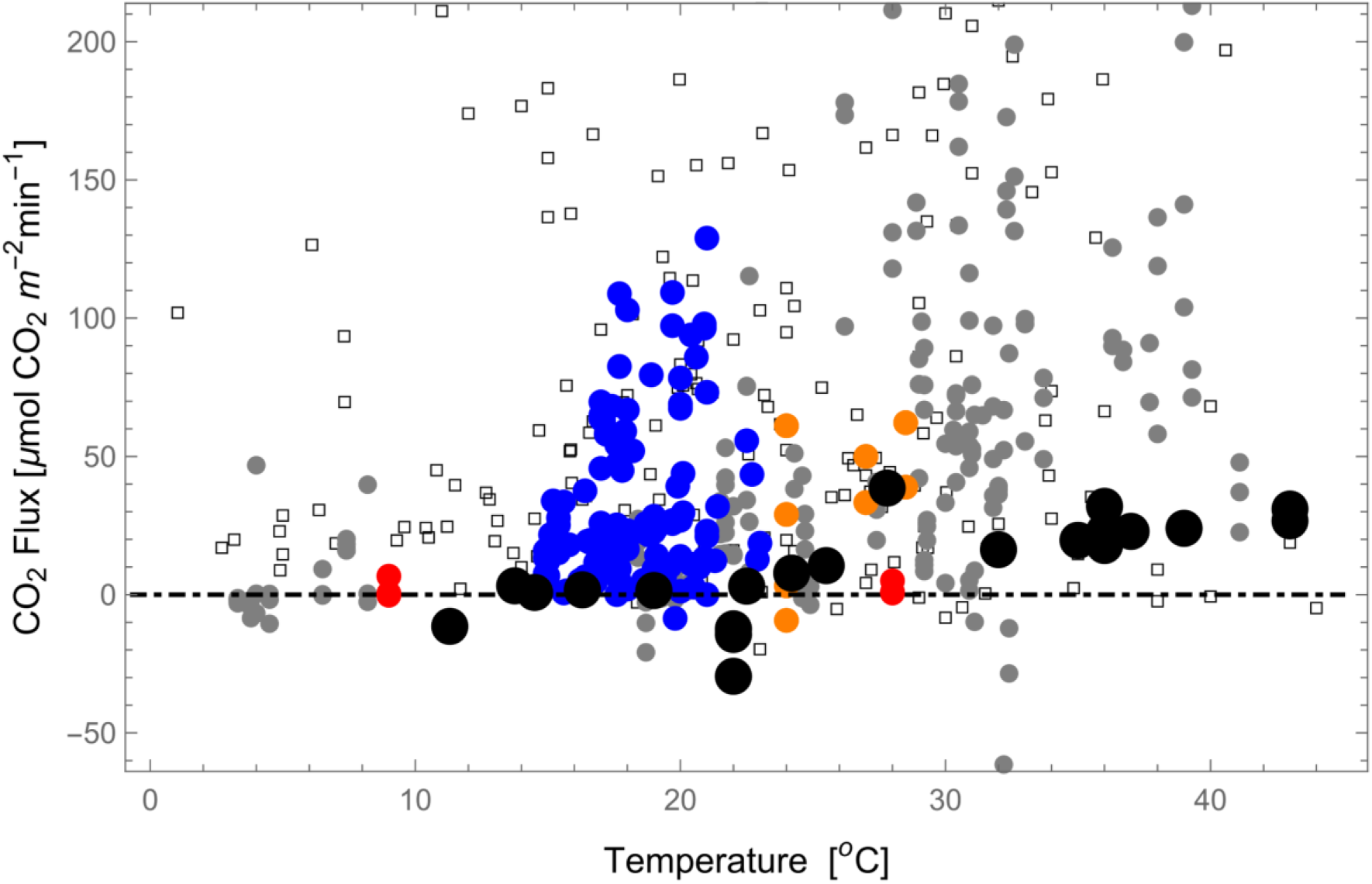
Relationship between CO_2_ fluxes and temperature reported in different hypersaline systems (dots) and dry freshwater systems (squares). CO_2_ fluxes reported in this plot were obtained using the closed chamber method. Black dots: this study; red dots: a hypersaline lake (Bourhane et al., 2023); orange dots: hypersaline tidal flats (Brown et al., 2021); grey dots: exposed Great Salt Lake bed sediments (Cobo et al., 2024); blue dots: hypersaline lake (Thomas et al., 2022); grey squares: exposed dry sediments from freshwater ecosystems (Keller et al., 2020). The range of “y” axes is constrained in the –50 to 200 μmol CO^2^ m^-2^ min^-1^ interval. Appendix 9 shows the same plot but with the full “y” range.

### 4g) CO_2_ fluxes in fractured OrgMinL

On September 16^th^, 2024, in Plot 2, the OrgMinL showed a network of fractures that typically formed five-sided polygons, where the sides measured 30–50 cm. The fractures had a width that exceptionally reached up to a centimetre, but they were usually smaller than 1.5 mm (Appendix 3) and in some cases they were often salt-sealed. In four out of six cases, the net CO_2_ flux was higher at the fractured sites than at the adjacent unfractured sites and, on average, net CO_2_ flux at fractured sites (54.6.3±26.3 μmol CO_2_/m^2^/min) was higher than at unfractured sites (43.4±19 μmol CO_2_/m^2^/min). However, the difference was not significant (t_paired_ =0.87, d.f.=5, p>0.05). Fracture density at the six selected points ranged between 6.7 and 11.5%. However, the net CO_2_ flux recorded at these six fractured locations was not significantly related to fracture density (r^2^=0.13, p>0.05).

### 4h) Ex-situ estimates. Light-dark incubations. The influence of photosynthesis on CO_2_ flux

*Exsitu* Exp#1): The samples used in this experiment had an a_w_ ranging between 70.5% and 83.94%, with a chlorophyll-a content in the OrgMinL averaging 1.3±0.26 μg cm^-**2**^. Net CO_2_ flux under light and dark conditions averaged 14.9±6.3 μmol min^-1^m^-2^ and 12±7.2 μmol min^-1^m^-2^ respectively. The difference was not significant (paired-t test, t=1.68, p>0.05, g.l.=11; Appendix 6).

*Exsitu* Exp #2): The a_w_ of OrgMinL samples averaged 80.5 ±1.3%, with a chlorophyll-a concentration identical to that measured in the previous experiment. The net CO_2_ emission fluxes averaged 8.4±6.4 CO_2_ μmol min^-1^m^-2^ and 7±2.2 CO_2_ μmol min^-1^m^-2^ during light and dark conditions respectively. The difference between averages was not significant (paired-t test, t=0.61, p>0.05, g.l.=7; Appendix 7).

*Ex-situ Exp #3):* The four samples that were submerged in water for a week showed a moderate increase in chlorophyll-a content up to 2.1±0.9 μg cm^-2^. In all measurements, CO_2_ release was higher in dark conditions. In two cases, under light conditions, a decrease in CO_2_ was observed (i.e., negative CO_2_ flux). The gross primary production in the samples averaged 5.5±0.8 CO_2_ μmol min^-1^m^-2^ (Appendix 8).

## 5) Discussion

At la Muerte, temperature and sediment water availability regulate CO_2_ flux at the sediment–atmosphere interface: emissions peak when temperature is high and sediment humidity low. This pattern matches findings reported in hypersaline systems (Cobo et al., 2024; Thomas et al., 2022; Alfadhel et al., 2024; Appendix 1) and in dry freshwater sediments (Keller et al., 2020). Therefore, this dependence of CO_2_ fluxes on temperature and humidity is probably a unifying characteristic of all continental aquatic systems subjected to severe water stress, regardless of their salinity.

Information on CO_2_ fluxes in hypersaline systems remains limited to a few case studies (Appendix 1). Focusing on closedchamber studies, our estimates fall within the same range as those reported for hypersaline tidal flats (Brown et al., 2021; Leopold et al., 2015) and for other hypersaline lakes located very close to la Muerte (La Salineta-Bourhane et al., 2023 - and Jabonera, Laguna Grande, and Laguna Pequeña-Thomas et al., 2022 - Figure 7). Net CO_2_ emission rates from la Muerte and Salineta are in the lower end of the range of estimates reported at the other three lakes. Under dry conditions, both Salineta and La Muerte form a surficial salt crust while such crusts have not been reported at Jabonera, Laguna Grande, and Laguna Pequeña. Further, the field tests revealed that the OrgMinL from La Muerte playa-lake, despite its thinness, significantly reduces CO_2_ fluxes across the sediment–atmosphere interface when the lake is dry. All together this suggests that the salt crust layer, when present, not only exerts physical control over water evaporation from underlying sediments (Fujimaki et al., 2006), but also influences the exchange of gases such as CO_2_.

Under dry conditions at la Muerte, the highest CO_2_ fluxes occurred beneath the OrgMinL, coinciding with peak water activity and organic-matter accumulation, indicating a microbial hotspot just below the organomineral layer. By contrast, although the OrgMinL hosts the highest bacterial density with the highest potential for organic-matter degradation (Boadella et al., 2024), its microbial activity is likely constrained by low water activity.

The OrgMinL shows domes, blisters and folds attributed to pressure exerted from water vapour and microbial-derived gases released from underlying sediments (Gerdes et al., 2007). Such formations are common in saline lakes (Noffke et al., 2001; Sanz-Montero and Rodríguez-Aranda, 2013) and has been reproduced in laboratory (Licsandru et al., 2022). The same gas pressure likely promotes their subsequent rupture and collapse. In la Muerte, lenticular-shaped, gypsum crystals are the main mineral precipitating in the underlying sediments whereas halite and other types of sulfates (mainly bloedite) predominate at the surface. Subsurface gypsum exhibits evidence of dissolution and persistence at depth, a pattern consistent with observations from other hypersaline lakes (Cabestrero et al., 2018). The chemical stability of gypsum suggests that this mineral is not involved in CO_2_ reactions.

Upon drying, the OrgMinL progressively loses its cohesiveness as microbes and their EPS degrade, serving as a locus for mineral precipitation (Del Buey et al., 2021). These minerals distribute anisotropically across and further harden the OrgMinL. Their shapes and arrangements produce a coherent, yet irregular crystalline assemblage with open porosity. In sum, minerals precipitating within the OrgMinL together with the extracellular polymers (EPS) help preserve its morphology and coherence (Mees and Tursina, 2018; Fulaz et al., 2019). In this context, an EPS compositional shift toward a higher proteintopolysaccharide ratio—proteins being effective adhesives (Santschi et al., 2020)—observed in the la Muerte biofilm has been interpreted as a drying signal (J. Boadella, unpublished data). Differences between surface and subsurface organomineral features further suggest distinct microbial communities in the OrgMinL and the underlying layers.

Overall, this thin, rigid upper layer, though not fully impermeable, can physically impede gas exchange at the atmosphere–sediment interface. Beyond this physical effect, a fraction of the heterotrophically produced CO_2_ in underlying sediments (Boadella et al., 2023) may be chemically retained. Authigenic carbonates detected within OrgMinL pores indicate that some CO_2_ binds to cations (Ca_2_^+^, Mg_2_^+^) adsorbed to organic matter and precipitates as calcite or dolomite (Chang et al., 2017).

Salt crust thicknesses in hypersaline systems span a few centimetres (Finstad et al., 2016) to over a meter (Bowen et al., 2018). Thus, if the CO_2_exchange impact observed at la Muerte lake arises under such a thin crust, stronger effects would be expected in systems with thicker salt crusts.

The OrgMinL at la Muerte hosts photosynthetic organisms, chiefly cyanobacteria (Berlanga et al., 2024; Boadella et al., 2024). Because field incubations used transparent chambers, photosynthetic primary producers could reduce CO_2_ emission rates (Leopold et al., 2015; Lovelock, 2008). Therefore, the reported increase in CO₂ emission after OrgMinL removal might be explained by the elimination of those producers inhabiting that layer. Nonetheless, the highest gas emissions after OrgMinL removal occurred during the dry period, when chlorophyll was highly degraded and at minimum concentrations, indicating scarce and largely inactive photosynthetic producers—consistent with microphotographs. Ex-situ Exp #1 and #2 likewise showed no clear evidence of substantial impact of photosynthesis on CO_2_ fluxes. Drying promotes salt precipitation and lowers water activity, suppressing the activity and diversity of primary producers in hypersaline habitats (Zhong et al., 2016; Yue et al., 2019; Lindsay et al., 2019). Thus, during dry periods, photosynthetic uptake of CO_2_ released by underlying sediments is likely minimal.

In contrast under water-saturated conditions, in-situ CO_2_ removal increased, coinciding with chlorophyll concentration in the OrgMinL nearly an order of magnitude higher than under dry conditions. Wetting dissolves salts and replaces the rigid crust with a slimy, glue-like biofilm that traps clastics and clay and grows under favorable conditions (Appendix 3). Indeed Ex-situ Exp #3 detected a photosynthetic effect on CO_2_ exchange only when chlorophyll was boosted in OrgMinL samples submerged for one week. Logically, the OrgMinL at la Muerte is not an impermeable barrier: gases can escape through cracks and pores. Although in-situ measurements did not show an unequivocal increase in CO_2_ flux at fractures, at four of six locations the highest fluxes coincided with cracked sediments, suggesting fractures may facilitate release. Additional escape pathways include abundant tunnels and burrows within the upper 20 cm. Deer and other animal footprints disrupt the OrgMinL and indirectly enhance CO_2_ efflux. Branched burrows and tunnels, together with piles of peloids attributed to insect excavations (Sanz-Montero et al., 2012), can raise sediment porosity by up to 34%. A wasp emerged from a tunnel (Appendix 3), and arthropod fragments occurred in samples. The presence, abundance, and depth of burrows likely facilitate gas exchange: they promote CO_2_ seepage from sediments to the atmosphere and the diffusion of atmospheric O_2_ into sediments, enhancing aerobic decomposition of stored organic matter and, consequently, CO_2_ release during the hottest, driest conditions (Figure 8).

**Fig. 8.**
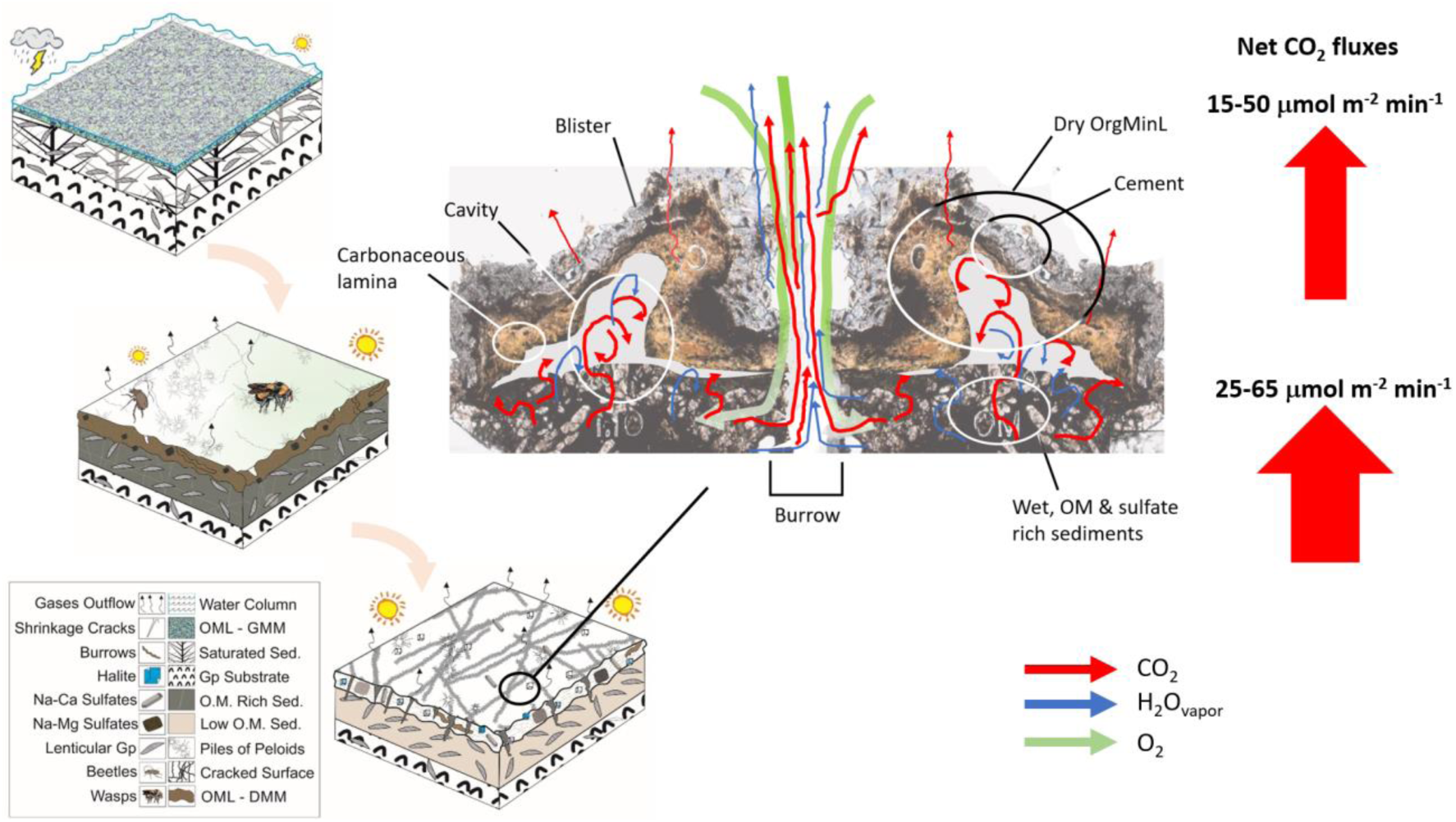
Model of organo-sedimentary evolution in La Muerte from wet to dry conditions, with the formation of the organic-mineral layer covering the sediment that limits CO_2_ fluxes (the red arrow) to the atmosphere during dry conditions.

Overall, as illustrated in Figure 7 (and Appendix 9), CO_2_ fluxes reported in hypersaline dry systems are typically lower than those reported in a global study focused on dry sediments from freshwater ecosystems (Keller et al., 2020). Therefore, endorheic dry hypersaline systems may not be a major source of CO_2_ when compared to freshwater (dry) systems. However, an important exception is Great Salt Lake, where exposed sediments emitted >100 µmol CO_2_ m⁻² min⁻¹ (Cobo et al., 2024 and Figure 7). Such emission rates are far above the rates recorded in the present study and the previous studies reported in Figure 7 (and also referred at the Appendix 1). Cobo et al. (2024) incubated in darkness, whereas our field incubations used natural light. One could expect higher emissions in darkness due to inhibited photosynthesis; however, our laboratory assays indicate that, in exposed dry OrgMinL from La Muerte, primary production is negligible relative to secondary production. Consistently, CO_2_ fluxes from hypersaline tidal flats show no large light–dark contrasts (Brown et al., 2021). Thus, light regime alone is unlikely to account for the discrepancy between Great Salt Lake and other studies. Instead, this divergence shows that our understanding of how the carbon cycle works in endorheic systems is still limited and highlights the need to broaden observations across sites and conditions.

## Conclusions

a. Our understanding of CO_2_ exchange in endorheic hypersaline shallow lakes during prolonged drought remains limited. Available data indicate that temperature, water availability, and sediment organic-matter content chiefly control flux magnitude, mirroring patterns reported for dry sediments in temporary freshwater ecosystems.
b. The structure and properties of the mineral-organic crust that forms and hardens at the surface during drying is fundamental not only to mitigate water evaporation but also to modulate CO_2_ exchange with the underlying sediments and the atmosphere.
c. Although subjected to extreme water stress, which inhibits biological activity in the organo-mineral surface layer, its mere presence helps to preserve biotic activity in the underlying moist sediments. Furthermore, its morphology is also likely indicative of transient gas accumulation.
d. The organomineral crust is not an impermeable gas barrier. Prolonged, severe drought compromises its integrity, generating cracks that, together with bioinduced burrows and tunnels, likely create preferential pathways for CO_2_ escape to the atmosphere. Simultaneously, these structures enhance oxygen diffusion into sediments, stimulating aerobic heterotrophy in underlying layers, and thereby increasing CO_2_ release.
e. Overall, this study highlights that a more comprehensive understanding of how hypersaline ecosystems exchange gases with the atmosphere needs to incorporate an analysis of the organic and mineralogical composition and structure of the crust that can form on their surface.

## Acknowledgments and funding

This study was funded by MCIN/AEI/10.13039/501100011033 (PID2021-123735OB-C22/C21). The authors thank the Instituto Aragonés de Gestión Ambiental (INAGA) for sampling authorization (permit INAGA 500201/24/2022/09146). We thank M. Millán Gómez and A. Peñaranda Martinez for performing the ex-situ experiments.

