## Supplemental material for "CO_2_ emission at a hypersaline shallow lake at sediment-atmosphere interface. The significance of the organomineral upper crust as an active barrier"

Berlanga M. Departament de Biologia, Sanitat i Mediambient, Secció de Microbiologia, Facultat de Farmàcia i Ciències de l'Alimentació, Universitat de Barcelona, Barcelona, Spain

Picart P. Departament de Biologia, Sanitat i Mediambient, Secció de Microbiologia, Facultat de Farmàcia i Ciències de l'Alimentació, Universitat de Barcelona, Barcelona, Spain

Garcia de Fuente Y. Institute of Aquatic Ecology, University of Girona, Spain; Department of Ecology and Hydrology, Faculty of Biology, University of Murcia, Murcia, Spain

Gomez R. Department of Ecology and Hydrology, Faculty of Biology, University of Murcia, Murcia, Spain

Urmeneta J. Departament de Genètica, Microbiologia i Estadística and Biodiversity Research Institute, Facultat de Biologia, Universitat de Barcelona, Barcelona, Spain

Romaní A. M. Institute of Aquatic Ecology, University of Girona, Spain

Esther Sanz-Montero. Department of Mineralogy and Petrology, Complutense University of Madrid, Madrid, Spain

**Appendix 1**

| System | $CO_2$ flux range (mmol $m^{-2}$ $h^{-1}$) | Method | Comments | Reference |
|---|---|---|---|---|
| Great Salt Lake (Utah, United States) | -2.5 - 41.8 | Incubation with opaque chamber | $CO_2$ net emission prevails under dry conditions | Cobo, Goldhammer, & Brothers. (2024). |
| Hypersaline tidal flats (Australia and Brazil) | -0.6 - 3.7 | Incubation with chamber under light and dark conditions | Higher $CO_2$ net emissions typically recorded in dark incubations. Higher $CO_2$ uptakes were recorded during dry season | Brown et al. (2021). |
| Hypersaline shallow lakes in Monegros (Spain) | -0.51 - 7.7 | Incubation with chambers | $CO_2$ net emission increases under dry conditions and with increasing temperature | Bourhane et al. (2023).<br>Thomas et al. (2022). |
| Fuente de Piedra Lake (Spain) | -11.4- -1.9 | Eddy covariance | $CO_2$ net emission prevails under dry conditions | Alfadhel et al. (2024) |
| Lakes on the Tibetan Plateau (China) | -1.16 - 0 | Eddy covariance | $CO_2$ net uptake prevails especially during cold no-icing periods. | Li, Wang, & Ma. (2024). |
| Saline lake Qinghai-Tibet Plateau (China) | -6.6 - 3.3 | Eddy covariance | $CO_2$ net uptake prevails especially during ice-covered winter periods | Li et al. (2022). |
| Saline lake Qinghai-Tibet Plateau (China) | -0.75 - 2 | Water-air interface diffusion model estimation | $CO_2$ net uptake prevails. Impact of seasons or dry/wet conditions were not studied. | Wang et al. (2022). |
| Bosten Lake (China) | 0.3 - 0.95 | Water-air interface diffusion model estimation | $CO_2$ net emission prevails. Increased salinity in water column stimulated $CO_2$ emission | Liao et al. (2024). |

Table SM1. Range of $CO_2$ fluxes reported in studies that measured $CO_2$ exchange in inland saline ecosystems. Negative $CO_2$ fluxes indicate net uptake; positive fluxes indicate net loss.

**Appendix 2**

The following plot shows the photosynthetic response of an OrgMinL sample submerged in water. Chlorophyll fluorescence measurements were performed with a Diving-Pam Chlorophyll fluorometer by Heinz Walz GmbH (Germany). The sensor was placed vertically (i.e. 90$^{o}$ geometry measurement) respect to the sample at a distance of 0.4 cm with a darkening adapter.

Two OrgMinL samples were used (5 x 5 cm) in this test. One was submerged in water during a week. The second stayed dry (i.e. control). The plot shows that the photosynthetic performance of the samples submerged in water rapidly increases during the first 48 hrs. and then stabilizes.

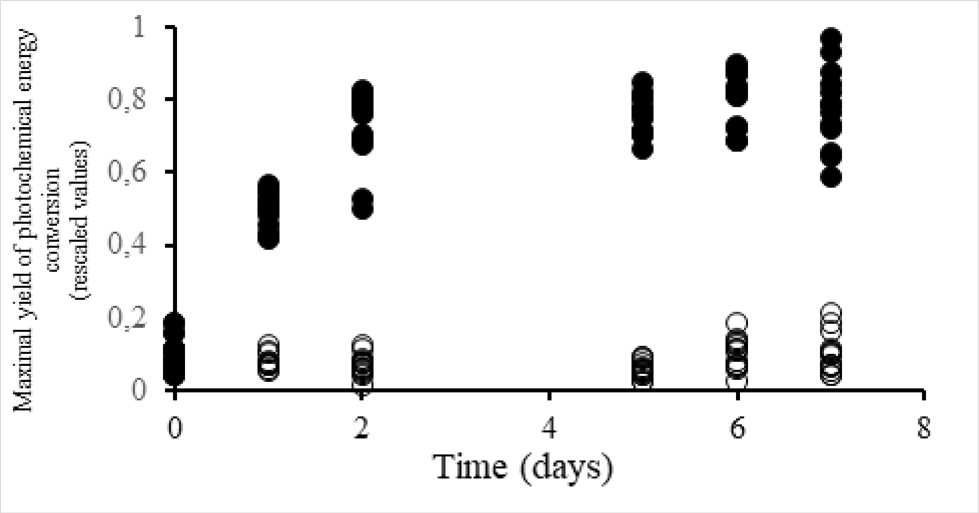


Figure SM2. Maximal yield of photochemical energy (a proxy photosynthetic efficiency) in a OrgMinL sample submerged in water during a week (black dots). Response is compared with that produced by a dry sample (open circles). Each dot is a single measurement.

**Appendix 3**

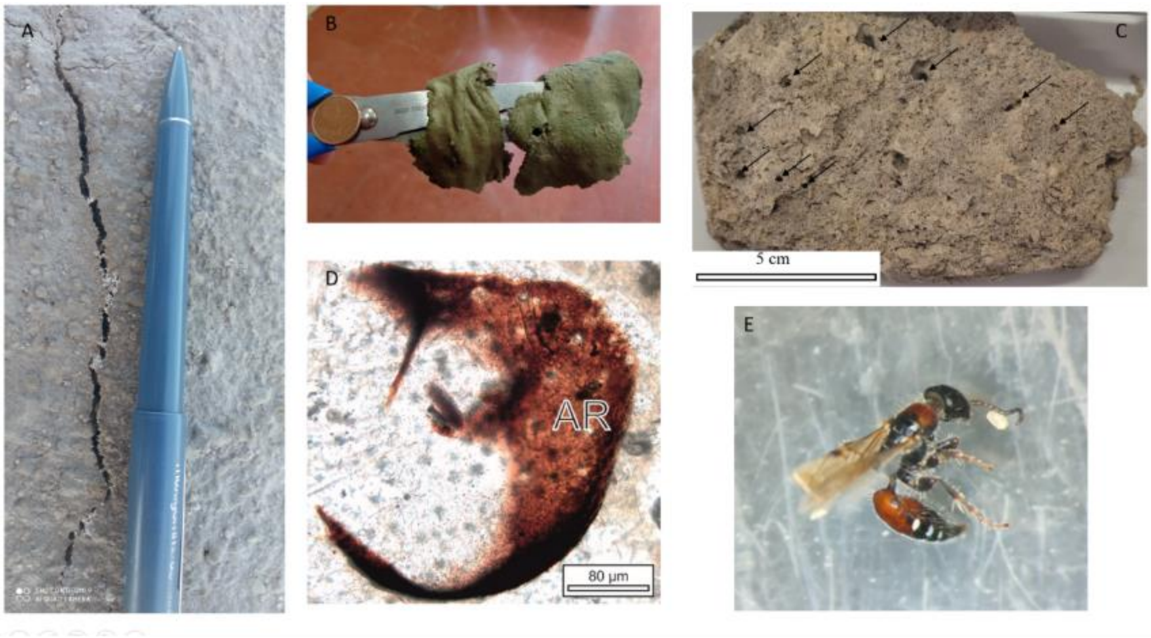


Figure SM3. A) Detail of a salt crust fracture observed at plot Area 2 on September 2024; B) detail of the wet upper slimy biofilm (picture taken on January 2024); C) Aspect of the dry sediment below the upper crust.
Black arrows indicate vertical burrows likely product of insect activities. D) microphotograph of a porous showing a fragment of an insect. E) a small wasp captured when was emerging from a burrow.

**Appendix 4**

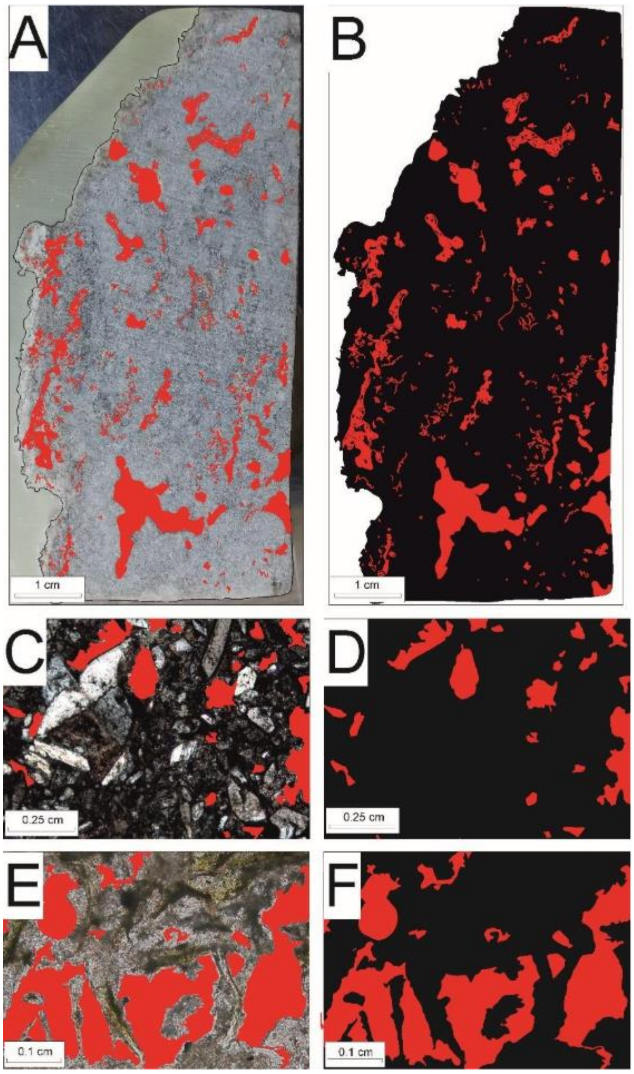


Figure SM4. Impregnated plug (A-B) and thin section photomicrographs under polarized light (C-F) for sediment samples. Areas delimited for porosity quantification are marked in red (A-F), while sediments are highlighted in black (B, D, and F). Notice that A-B allow for the measurement of larger porosity on a smaller scale, whereas C-D show the smaller porosity within the sediment beneath the crust, organomineral layer, depicted in E-F.

**Appendix 5**

In situ $CO_2$ fluxes in the upper 20 cm of sediments.

The following table shows the environmental conditions and content of water, organic matter, bacterial density and net $CO_2$ flux in a sediment core 20 cm depth. Measurements performed July 30th 2024 under dry conditions.

| Z (cm) | T ($^oC$) | $a_W$ (%) | WatCon (%) | AFDW (%) | BactDens ($10^6$ $g^{-1}$) | Bglu (nM MUF $cm^{-3}$ $h^{-1}$) | $CO_2$ Flux (µmol $CO_2$ $m^{-2}$ $min^{-1}$) |
|---|---|---|---|---|---|---|---|
| 0 | 41 | 74.0 | 18.94 | 14.85 | 23.85 | 11.13 | 34.8 |
| 0.3 | 41 | 73.8 | 16.44 | 20.93 | 22.11 | 13.26 | 45.1 |
| 1 | 39 | 80.6 | 17.96 | 19.45 | 22.06 | 6.28 | 29.8 |
| 3.5 | 38 | 80.8 | 17.52 | 19.07 | 21.43 | 9.67 | 52.6 |
| 8.5 | 36 | 82.1 | 18.58 | 18.91 | 20.39 | 1.19 | 33.6 |
| 14.2 | 36 | 86.4 | 19.34 | 17.4 | 20.57 | 3.55 | 13 |
| 20 | 36 | 86.0 | 19.27 | 17.44 | 18.7 | 1.32 | 7.1 |

Table SM5a. Environmental conditions, bacterial density, exoenzymatic activity values and $CO_2$ fluxes recorded in the upper 20 cm of the sediment.

$a_W$: Water activity; WatCon: Water content in sediments; AFDW: ash free in sediments (a proxy of particulate organic matter content): BactDens: bacterial density; Bglu: Beta glucosidase exoenzymatic activity (a proxy of potential ability of microbiota to degrade labile organic matter.

According to these results, the best model (that with the lowest BIC value) is that related the Net $CO_2$ fluxes with AFDW and $a_W$ according to the following multiple linear model (Net$CO_2$ flux = -4,4+5.03 AFDW+4.53 $a_W$, $r^2$=0.8, $p<0.05$, BIC=1.8, the Anova table is the table SM4b).

| Descriptors | d.f. | MSE | F-statistic | P |
|---|---|---|---|---|
| $a_W$ | 1 | 0.45 | 11.89 | 0.026 |
| AFDW | 1 | 0.16 | 4.119 | 0.112 |
| Error | 4 | 0.04 | | |
| Total | 6 | | | |

Table SM5b. The Anova table of the multiple linear model. More details at section 4e. d.f. : degree of freedom: MSE: Means Square error

### Appendix 6

*Ex-situ Exp #1:* Day-night incubations with undisturbed sediment samples including the OrgMinL and 4-7 cm of the underlying sediments.

Results: The samples used in this experiment had an $a_w$ ranging between 70.5% and 83.94%, with a chlorophyll-a content in the OrgMinL averaging 1.3±0.26 µg cm$^{-2}$.

Net $CO_2$ flux under light conditions averaged 14.9±6.3 µmol min$^{-1}$m$^{-2}$ and was slightly higher than that recorded in dark conditions (12±7.2 µmol min$^{-1}$m$^{-2}$). However, the difference was not significant (paired-t test, t=1.68, p>0.05, g.l.=11) (Table SM6).

Ex-situ Exp#1. Results

| Sample ID + Replicate | Treatment | $a_w$ (%) | Temp (°C) | $CO_2$ Flux (µmol m$^{-2}$ min$^{-1}$) | $r^2$ |
|---|---|---|---|---|---|
| *#1_R1* | Light | 83.94 | 25.83 | 12.3 | 0.81 |
| | Dark | | 25.79 | 11.92 | 0.82 |
| *#1_R2* | Light | | 26.15 | 10.19 | 0.97 |
| | Dark | | 26.6 | 8.14 | 0.84 |
| *#1_R3* | Light | | 35.72 | 23.7 | 0.97 |
| | Dark | | 31.98 | 18.69 | 0.92 |
| *#2_R1* | Light | 81.7 | 22.63 | 8.56 | 0.86 |
| | Dark | | 21.62 | 2.61 | 0.29 |
| *#2_R2* | Light | | 23.44 | 14.53 | 0.81 |
| | Dark | | 23.57 | 5.97 | 0.64 |
| *#2_R3* | Light | | 32.97 | 23.15 | 0.97 |
| | Dark | | 30.04 | 30.43 | 0.97 |
| *#3_R1* | Light | 83.84 | 23.67 | 12.26 | 0.93 |
| | Dark | | 25.36 | 7.85 | 0.93 |
| *#3_R2* | Light | | 29.16 | 4.54 | 0.81 |
| | Dark | | 31.55 | 11.42 | 0.96 |
| *#3_R3* | Light | | 32.22 | 10.6 | 0.98 |
| | Dark | | 28.73 | 13.77 | 0.95 |
| *#4_R1* | Light | 70.5 | 28.74 | 15.3 | 0.91 |
| | Dark | | 28.57 | 7.05 | 0.49 |
| *#4_R2* | Light | | 28.56 | 22.09 | 0.91 |
| | Dark | | 28.99 | 14.74 | 0.74 |
| *#4_R3* | Light | | 29.1 | 21.52 | 0.86 |
| | Dark | | 29.14 | 11.36 | 0.64 |

Table SM6. Results of Ex-situ Exp#1. More details at section 4g. Significance levels of all estimates are p<0.001.

**Appendix 7**

*Ex-situ Exp #2:* Day-night incubations with undisturbed OrgMinL samples (without the underlying sediments)

Results: the $a_w$ of OrgMinL samples averaged 80.5 ±1.3%, with a chlorophyll-a concentration identical to that measured in the previous experiment. A rise in $CO_2$ concentration was consistently observed whether the samples were incubated in light or dark conditions, further indicating that, under dry conditions, the OrgMinL is a net $CO_2$ emitter regardless of light conditions. Overall, $CO_2$ emission fluxes averaged 8.4±6.4 $CO_2$ μmol $min^{-1}m^{-2}$ and 7±2.2 $CO_2$ μmol $min^{-1}m^{-2}$ during light and dark conditions respectively. The difference between averages was not significant (paired-t test, t=0.61, p>0.05, g.l.=7). Overall, under dry conditions, primary production in the OrgMinL seemed to be negligible (Table SM7).

Results of Ex-situ Exp#2:

| Sample ID | Treatment | Temp (°C) | $CO_2$ Flux ($CO_2$ μmol $m^{-2}$ $min^{-1}$) | $r^2$ |
|---|---|---|---|---|
| #1 | Light | 25.10 | 7.12 | 0.93 |
| | Dark | 26.30 | 6.65 | 0.98 |
| #2 | Light | 26.56 | 7.93 | 0.96 |
| | Dark | 27.46 | 8.98 | 0.99 |
| #3 | Light | 20.89 | 6.04 | 0.96 |
| | Dark | 21.92 | 10.40 | 0.99 |
| #4 | Light | 24.59 | 6.47 | 0.96 |
| | Dark | 25.58 | 8.18 | 0.99 |
| #5 | Light | 22.95 | 2.20 | 0.81 |
| | Dark | 23.56 | 4.17 | 0.97 |
| #6 | Light | 24.96 | 3.27 | 0.95 |
| | Dark | 24.97 | 4.14 | 0.94 |
| #7 | Light | 34.68 | 11.78 | 0.99 |
| | Dark | 31.64 | 7.34 | 0.94 |
| #8 | Light | 35.41 | 22.63 | 0.99 |
| | Dark | 31.30 | 6.38 | 0.88 |

Table SM7. Results of Ex-situ Exp#2. More details at section 4g. Significance levels of all estimates are $p<0.001$.

**Appendix 8**

*Ex-situ Exp #3:* Day-night incubations with undisturbed OrgMinL samples submerged in aerated water during a week.

Results:

| Sample ID | Treatment | Temp ($^oC$) | $CO_2$ Flux ($CO_2$ μmolm$^{-2}$ min$^{-1}$) | $r^2$ |
|---|---|---|---|---|
| #1 | Light | 22.29 | 1.14 | 0.66 |
| | Dark | 23.82 | 5.93 | 0.95 |
| #2 | Light | 20.44 | 11.18 | 0.98 |
| | Dark | 21.98 | 16.14 | 0.96 |
| #3 | Light | 30.48 | -4.17 | 0.95 |
| | Dark | 28.01 | 1.87 | 0.79 |
| #4 | Light | 34.07 | -8.01 | 0.99 |
| | Dark | 27.19 | -1.59 | 0.74 |

Table SM8. Results of Ex-situ Exp#3. More details at section 4g. Significance levels of all estimates are $p<0.001$.

**Appendix 9.**

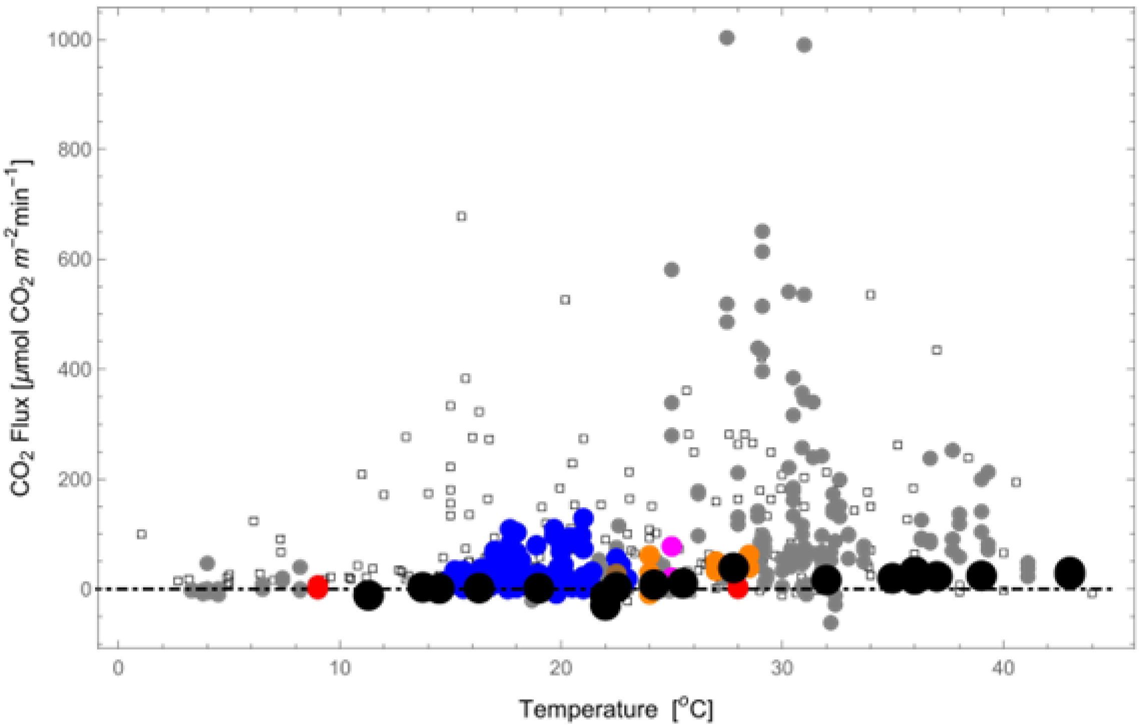


Figure SM9. This plot is the same as Figure 7 in the article but with the full "y" range. The Net $CO_2$ flux values reported in the present study (black dots) fall within the lower range of reported literature values. Symbols are the same as figure 7. References are in Figure 7.
